# Elevated plasma ceramide levels in post-menopausal women

**DOI:** 10.1101/365304

**Authors:** Valentina Vozella, Abdul Basit, Fabrizio Piras, Natalia Realini, Andrea Armirotti, Paola Bossù, Francesca Assogna, Stefano L. Sensi, Gianfranco Spalletta, Daniele Piomelli

## Abstract

Circulating ceramide levels are abnormally elevated in age-dependent pathologies such as cardiovascular diseases, obesity and Alzheimer’s disease. Nevertheless, the potential impact of age on plasma ceramide levels has not yet been systematically examined. In the present study, we quantified a focused panel of plasma ceramides and dihydroceramides in a cohort of 164 subjects (84 women) 19 to 80 years of age. After adjusting for potential confounders, multivariable linear regression analysis revealed a positive association between age and ceramide (d18:1/24:0) (β (SE) = 5.67 (2.38); *p* = .0198) and ceramide (d18:1/24:1) (β (SE) = 2.88 (.61); *p* < .001) in women, and between age and ceramide (d18:1/24:1) in men (β (SE) = 1.86 (.77); *p* = .0179). In women of all ages, but not men, plasma ceramide (d18:1/24:1) was negatively correlated with plasma estradiol (r = −0.294; *p* = .007). Finally, *in vitro* experiments in human cancer cells expressing estrogen receptors showed that incubation with estradiol (10 nM, 24 h) significantly decreased ceramide accumulation. Together, the results suggest that aging is associated with an increase in circulating ceramide levels, which in post-menopausal women may be at least partially dependent on lower estradiol levels.

## Introduction

The ceramides are key lipid constituents of mammalian cells. They regulate the structural properties of the lipid bilayer [1] along with its interaction with cellular proteins [2], and control many signalling processes, including cell survival [3], growth and proliferation [4], differentiation [5], senescence [6] and apoptosis [7,8]. Dysfunctions in ceramide-mediated signalling may contribute to the initiation and progression of a variety of age-dependent diseases. Human studies have shown the existence of abnormal plasma levels of various ceramide species – including ceramide (d18:1/18:0), (d18:1/22:0), (d18:1/24:0) and (d18:1/24:1) – in several conditions such as obesity [9], type-2 diabetes [10], hypertension [11], atherosclerosis [12] and other cardiovascular diseases [13]. Furthermore, elevated serum levels of long-chain ceramides have been linked to the increased risk of memory deficits [14] and may be predictive of hippocampal volume loss and cognitive decline in patients affected by Mild Cognitive Impairment [15]. Other studies have reported the existence of sex-dependent differences in circulating ceramides, albeit with apparently contrasting results [16-18]. For example, in a study of a large cohort of Mexican-Americans of median age 35.7 years, plasma ceramides were found to be higher in men than in women [17]. By contrast, in the Baltimore Longitudinal Study of Aging, whose participants were aged 55 or older, plasma ceramide concentrations were shown to be higher in women than in men [18].

Aging in rats and mice is associated with sexually dimorphic changes in the sphingolipid composition of several brain structures, including the hippocampus [19]. Age-dependent sphingolipid alterations have also been documented in peripheral rodent tissues [20]. In the present study, we hypothesized that aging in humans might be similarly associated with changes in ceramide levels. To test this idea, we profiled six ceramide and dihydroceramide species in lipid extracts of plasma from 164 subjects (84 women) of age 19 to 80 years, using liquid chromatography/mass spectrometry (LC-MS/MS). A primary criterion for subject selection was the absence of major medical illnesses and, particularly, of conditions that had been previously linked to ceramide alterations.

## Materials and Methods

### Study subjects

We recruited 164 Italian subjects (84 women) from 19 to 80 years of age (Table 1). There were no significant differences between male and female subjects with regard to age, years of education, ethnic background, cognitive status, as assessed by the Mini-Mental State Examination (MMSE), obesity and diabetes, hypertension and use of anti-hypercholesterolemic drugs. By contrast, there was difference in smoking status (20.23% women versus 7.5% men). Moreover, 6 of 44 pre-menopause women used contraceptives at time of enrollment. None of the post-menopause women was under hormone replacement therapy (HRT).

**Table 1.**
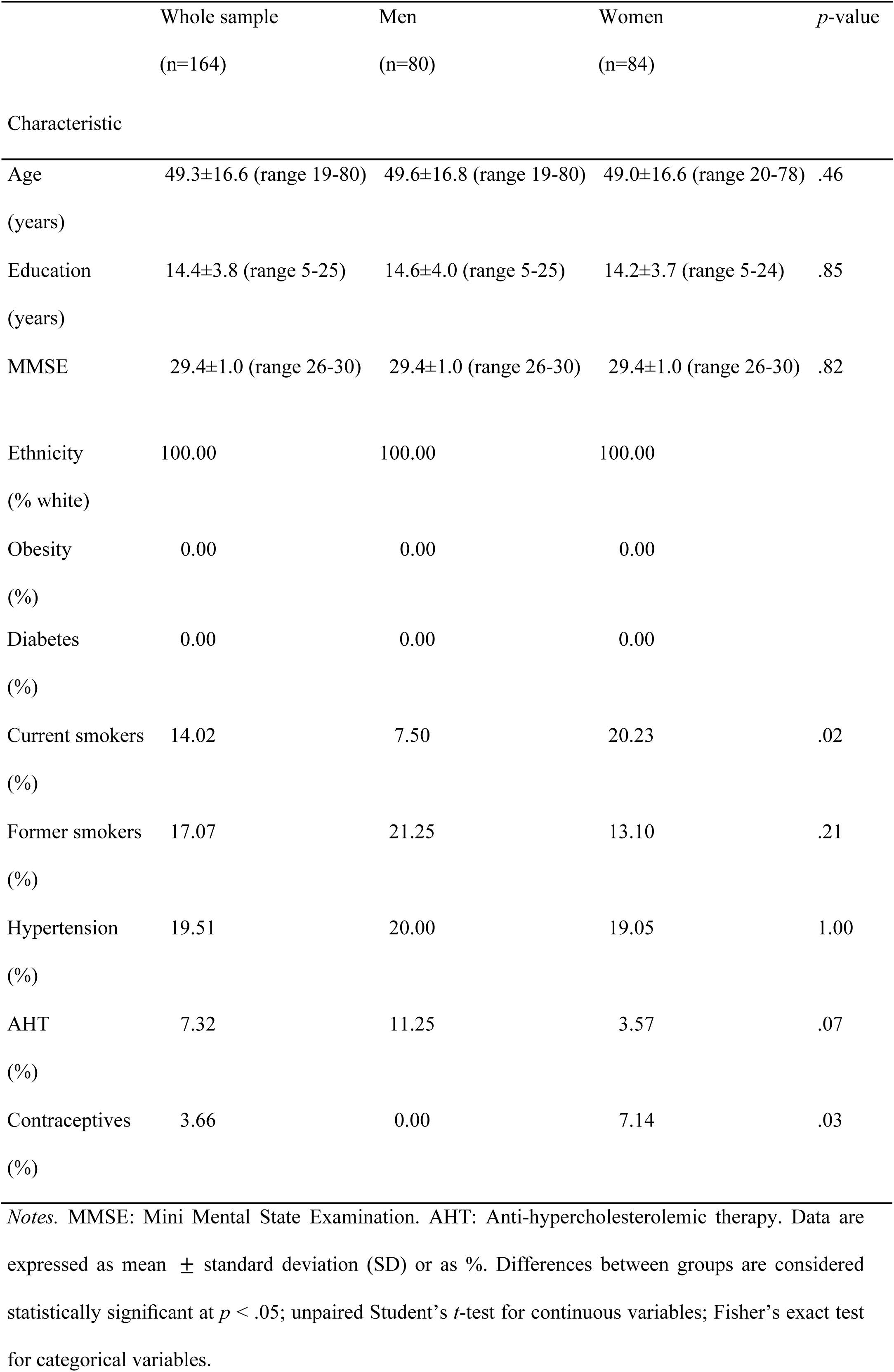
Sociodemographic and clinical characteristics of men and women included in the study.

Exclusion criteria were: (i) suspicion of cognitive impairment or dementia based on MMSE [21] (score ≤ 26, consistent with normative data collected in the Italian population) and confirmed by a detailed neuropsychological evaluation using the Mental Deterioration Battery [22] and clinical criteria for Alzheimer’s dementia [23] or Mild Cognitive Impairment [24]; (ii) subjective complaints of memory difficulties or other cognitive deficits, regardless of whether or not these interfered with daily life; (iii) vision and hearing loss that could potentially influence testing results; (iv) major medical illnesses (i.e., unstable diabetes; obesity; obstructive pulmonary disease or asthma; hematological and oncological disorders; pernicious anemia; significant gastrointestinal, renal, hepatic, endocrine, or cardiovascular system diseases; recently treated hypothyroidism); (v) current or reported psychiatric disease, as assessed by the Structured Clinical Interview for Diagnostic and Statistical Manual of Mental Disorders, 4th Edition, Text Revision (DSM-IV-TR SCID) [25] or neurological disease, as assessed by clinical evaluation; and (vi) known or suspected history of alcoholism or drug addiction. Finally, because ceramides have been previously involved in the pathogenesis of neurodegenerative disorders [15,26,27], we excluded subjects who showed brain abnormalities or vascular lesions as determined by using a recently published semi-automated method [28]. The menopausal status was prospectively assessed during clinical interviews. Women were defined as post-menopausal when showing 12 consecutive months of amenorrhea that was not due to other obvious pathological or physiological causes.

Blood collection and analyses were approved by the Santa Lucia Foundation Ethics Committee and complied with the ethical principles set out in the Declaration of Helsinki. A written consent form was signed by all participants after they received a full explanation of the study procedures.

### Variables examined in relation to ceramide levels

All variables were assessed for each patient using the same methods. Demographic variables considered in linear regression analysis included age and sex. Medical history covariates included hypertension, use of anti-hypercholesterolemic agents and use of contraceptives. Current and former smoking status was ascertained by an oral interview.

### Chemicals

Solvents and chemicals were purchased from Sigma Aldrich (Milan, Italy). Ceramide standards were from Avanti Polar Lipids (Alabaster, AL, USA).

### Blood collection

Blood was drawn by venipuncture in the morning after an overnight fast, and collected into 10-ml tubes containing spray-coated EDTA (EDTA Vacutainer, BD Biosciences, San Diego, CA, USA). Plasma was obtained by blood centrifugation at 400 × *g* at 4 °C for 15 min. The plasma divided into aliquots was stored at –80 °C until analyses.

### Lipid extraction

Lipids were extracted using a modified Bligh and Dyer method [29]. Briefly, plasma samples (50 μL) or cell pellets were transferred to glass vials and liquid-liquid extraction was carried out using 2 mL of a methanol/chloroform mixture (2:1 vol/vol) containing the odd-chain saturated ceramide (d18:1/17:0) as an internal standard. After mixing for 30 s, lipids were extracted with chloroform (0.5 mL), and extracts were washed with liquid chromatography-grade water (0.5 mL), mixing after each addition. The samples were centrifuged for 15 min at 3500 × *g* at room temperature. After centrifugation, the organic phases were collected and transferred to a new set of glass vials. To increase overall recovery, the aqueous fractions were extracted again with chloroform (1 mL). The two organic phases were pooled, dried under a stream of N_2_ and residues were dissolved in methanol/chloroform (9:1 vol/vol, 0.07 mL). After mixing (30 s) and centrifugation (10 min, 5000 × *g*, room temperature) the samples were transferred to glass vials for analyses.

### Ceramide quantification

Ceramides were analyzed by LC-MS/MS using an Acquity^®^ ultra-performance liquid chromatography (UPLC) system coupled to a Xevo triple quadrupole mass spectrometer interfaced with electrospray ionization (Waters, Milford, MA, USA), as previously described [29]. Lipids were separated on a Waters Acquity^®^ BEH C18 column (2.1 × 50 mm, 1.7 μm particle size) at 60 °C and eluted at a flow rate of 0.4 mL/min. The mobile phase consisted of 0.1% formic acid in acetonitrile/water (20:80 vol/vol) as solvent A and 0.1% formic acid in acetonitrile/isopropanol (20:80 vol/vol) as solvent B. A gradient program was used: 0.0–1.0 min 30% B, 1.0–2.5 min 30 to 70% B, 2.5–4.0 min 70 to 80% B, 4.0–5.0 min 80% B, 5.0–6.5 min 80 to 90% B, and 6.6–7.5 min 100% B. The column was reconditioned to 30% B for 1.4 min. The injection volume was 3 μL. Detection was done in the positive electrospray ionization mode. Capillary voltage was 3.5 kV and cone voltage was 25 V. The source and desolvation temperatures were set at 120 °C and 600 °C respectively. Desolvation gas and cone gas (N_2_) flow were 800 L/h and 20 L/h, respectively. Plasma and cell-derived ceramides were identified by comparison of their LC retention times and MS/MS fragmentation patterns with those of authentic standards (Avanti Polar Lipids). Extracted ion chromatograms were used to identify and quantify the following ceramides and dihydroceramides (d18:1/16:0) (*m/z* 520.3 > 264.3), (d18:1/18:0) (*m/z* = 548.3 > 264.3), (d18:1/24:0) (*m/z* = 632.3 > 264.3), (d18:1/24:1) (*m/z* = 630.3 > 264.3), (d18:0/24:0) (*m/z* = 652.5 > 634.5) and (d18:0/24:1) (*m/z* = 650.5 > 632.5). Data were acquired by the MassLynx software and quantified using the TargetLynx software (Waters, Milford, MA, USA).

### Estradiol quantification

Plasma 17-β-estradiol (E2) levels were quantified using a competitive binding immunoassay kit (Human E2 ELISA kit, Invitrogen, Milan, Italy) following manufacturer’s instructions. Briefly, plasma samples, controls and standard curve samples (50 μL) were incubated with E2-horseradish peroxidase conjugate (50 μL) and anti-estradiol antibody (50 μL) in a 96-well plate for 2 h, at room temperature, on a shaker set at 700 ± 100 rpm. Washing was carried out by completely aspirating the liquid, filling the wells with diluted wash buffer (0.4 mL) provided in the kit and then aspirating again. After repeating this procedure 4 times, chromogen solution (200 μL) was added to each well; reactions were run for 15 min and stopped adding 50 μL of the stop solution provided in the kit. Absorbance was measured at 450 nm and estradiol concentrations were calculated by interpolation from the reference curve.

### Cell cultures and treatment

The MCF7 human breast cancer cell line [30,31] was a kind gift of Dr. Gennaro Colella (Mario Negri Institute, Milan, Italy). Cells were cultured in phenol red-free Dulbecco’s Modified Eagle’s Medium (DMEM) (Gibco by Life Technologies, Carlsbad, CA, USA) supplemented with charcoal-stripped fetal bovine serum (10%) (Sigma Aldrich, Milan, Italy) to starve cells from steroid hormones (starvation medium), L-glutamine (2 mM), and penicillin/streptomycin (100 μg/mL), in a humidified atmosphere (5% CO_2_, 37 °C). Cells were seeded in 6-well plates (3 × 10^5^ cells/well) and cultured for 24 h. Estradiol (Sigma Aldrich, Milan, Italy) was dissolved in dimethyl sulfoxide (DMSO) and diluted in phenol red-free DMEM to a final concentration of 10 nM (0.1% final DMSO concentration). After 24 h incubation, the media were removed, cells were washed with phosphate-buffered saline, scraped and centrifuged (800 × *g*, 4 °C, 10 min). Protein concentrations were measured using the bicinchoninic acid assay (Pierce, Rockford, IL, USA) and cell pellets were stored at −80 °C until analyses.

### Statistical analyses

Results are expressed as mean ± SEM (standard error of the mean). Sex differences in baseline demographic and health-related characteristics were examined using Fisher’s exact test for categorical variables and unpaired Student’s *t*-test for continuous variables. Data were analyzed by unpaired Student’s *t*-test or 2-way ANOVA followed by Bonferroni post-hoc test. Pearson’s correlation coefficient assessed the pairwise correlation between estradiol and ceramide levels. Significant outliers were excluded using the Grubbs’ test. Multivariable linear regression method was used to assess the association between ceramides and patients’ characteristics. Differences between groups were considered statistically significant at values of *p* < .05. The GraphPad Prism software (GraphPad Software, Inc., La Jolla, CA, USA) and SAS 9.4 (SAS Institute, Cary, NC, USA) were used for statistical analyses.

## Results

### Plasma ceramide levels are positively correlated with age

The scatter plot reported in Fig 1A illustrates the total ceramide levels in plasma of individual women aged 20-78 years. Pearson’s analysis of the data revealed a statistically significant positive correlation between ceramide levels and age (r = 0.378; *p* = .0004). Because the largest accrual in plasma ceramides occurred between the age of 40 and 50 years, which is coincident with menopause, in a secondary analysis we grouped the data according to the subjects’ menopausal status. We found a statistically detectable difference between pre-menopausal women (20-54 years) and post-menopausal women (47-78 years) (Fig 1B). In particular, the levels of long-chain ceramide (d18:1/18:0) (*p* = .0035, unpaired Student’s *t*-test), very long-chain ceramides (d18:1/24:0) (*p* = .0012) and (d18:1/24:1) (*p* < .0001), and dihydroceramide (d18:0/24:1) (*p* = .0340) were higher in post-menopausal relative to pre-menopausal women (Fig 1B).

**Fig 1.**
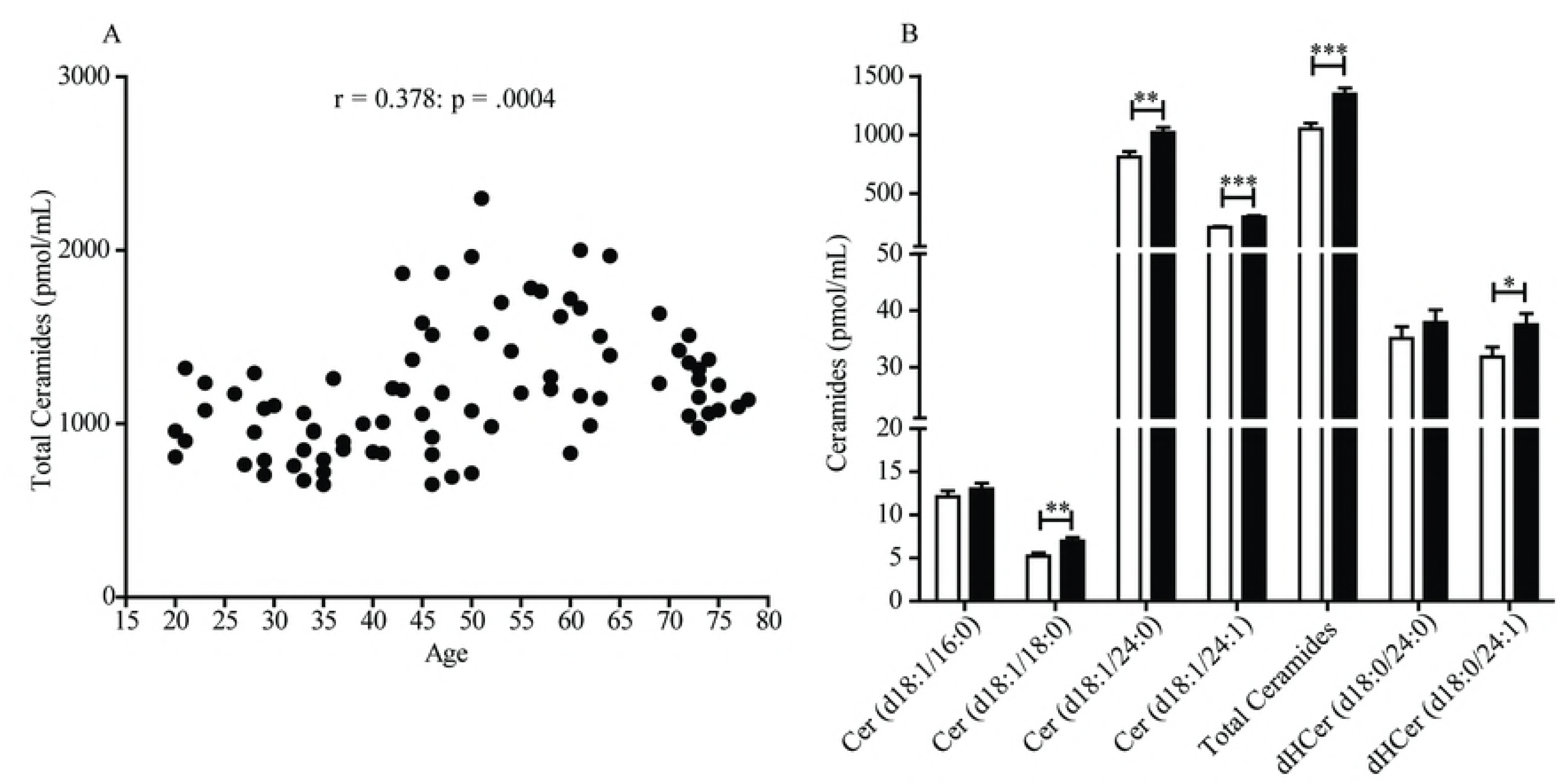
Scatter plot of plasma ceramide concentrations in women aged 20 to 78 years. (A) Total ceramide levels in 84 female subjects included in the study. Pearson’s correlation is considered statistically significant at *p* < .05. (B) Average levels of individual ceramide species in pre-menopausal women (20-54 years, n = 44, open bars) and post-menopausal women (47-78 years, n = 40, closed bars). Results are expressed as mean ± SEM. ^∗^*p* < .05, ^∗∗^*p* < .01, ^∗∗∗^*p* < .001; unpaired Student’s *t*-test.

No differences were found in the levels of ceramide (d18:1/16:0) (*p* = .3526) and dihydroceramide (d18:0/24:0) (*p* = .3633). In contrast with these findings in women, men showed no significant age-dependent increases in plasma ceramides (r = 0.143; *p* = .208) (Fig 2A). Male subjects in the age groups 19-54 and 55-80 years displayed comparable levels of circulating ceramide (d18:1/18:0) (*p* = .7112), (d18:1/24:0) (*p* = .7895), (d18:1/24:1) (*p* = .0847) and dihydroceramide (d18:0/24:1) (*p* = .9014). However, dihydroceramide (d18:0/24:0) was significantly lower in men >55 years, compared to younger men (*p* = .0003) (Fig 2B).

**Fig 2.**
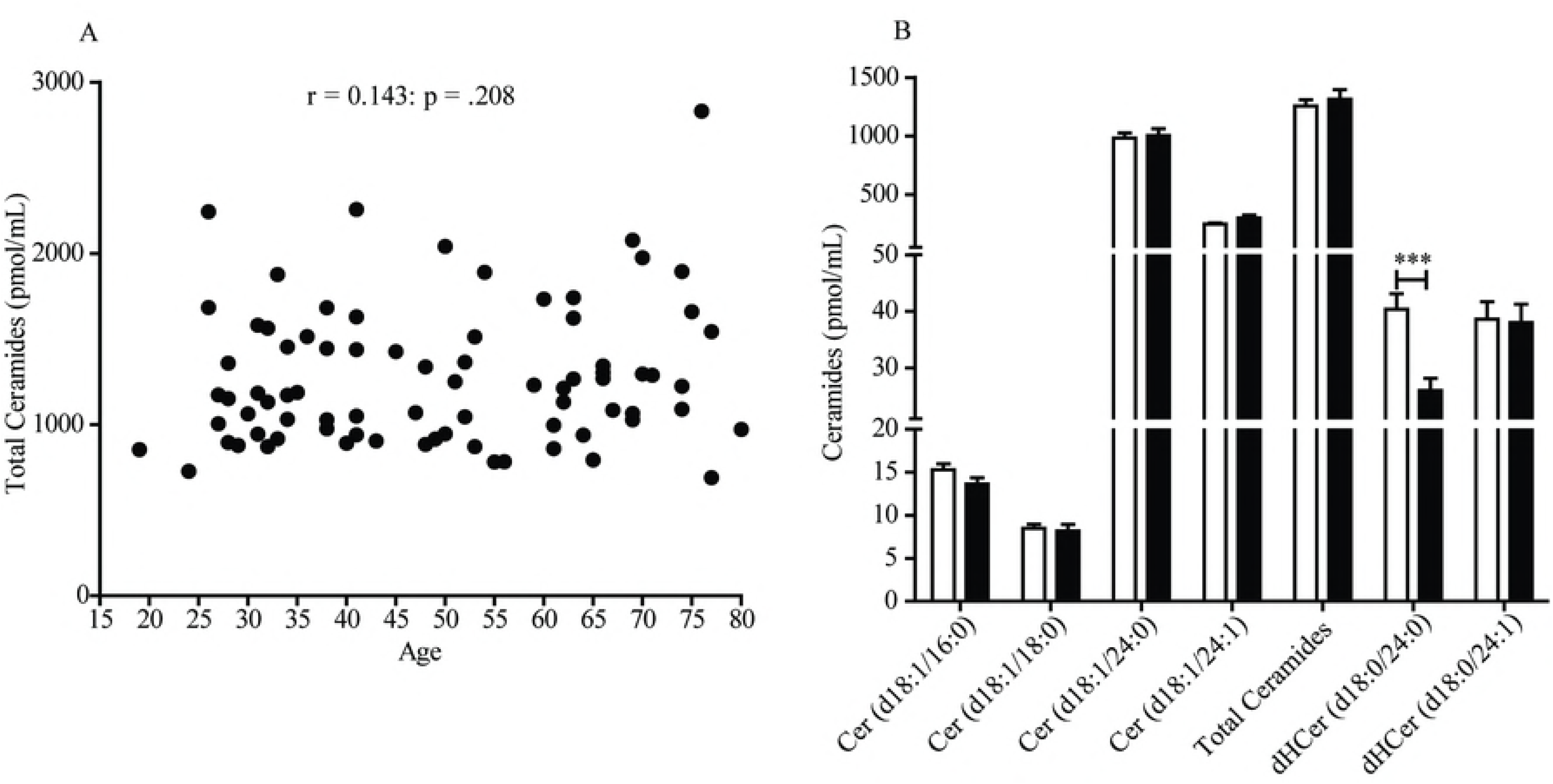
Scatter plot of plasma ceramide concentrations in men aged 19 to 80 years. (A) Total ceramide levels in 80 male subjects included in the study. Pearson’s correlation is considered statistically significant at *p* < .05. (B) Average levels of individual ceramide species in men aged 19-54 years (n = 48, open bars) and 55-80 years (n = 32, closed bars). Results are expressed as mean ± SEM. ^∗^*p* < .05, ^∗∗^*p* < .01, ^∗∗∗^*p* < .001; unpaired Student’s *t*-test.

In an additional analysis, we compared total ceramide levels in plasma of pre- and post-menopausal women with those measured in age-matched men (Fig 3). The results show that pre-menopausal women had significantly lower levels of circulating ceramides (*p* < .05, 2-way ANOVA followed by Bonferroni post-hoc test) relative to men of the same age (Fig 3A). The difference disappeared after menopause (*p* > .05) (Fig 3A).

**Fig 3.**
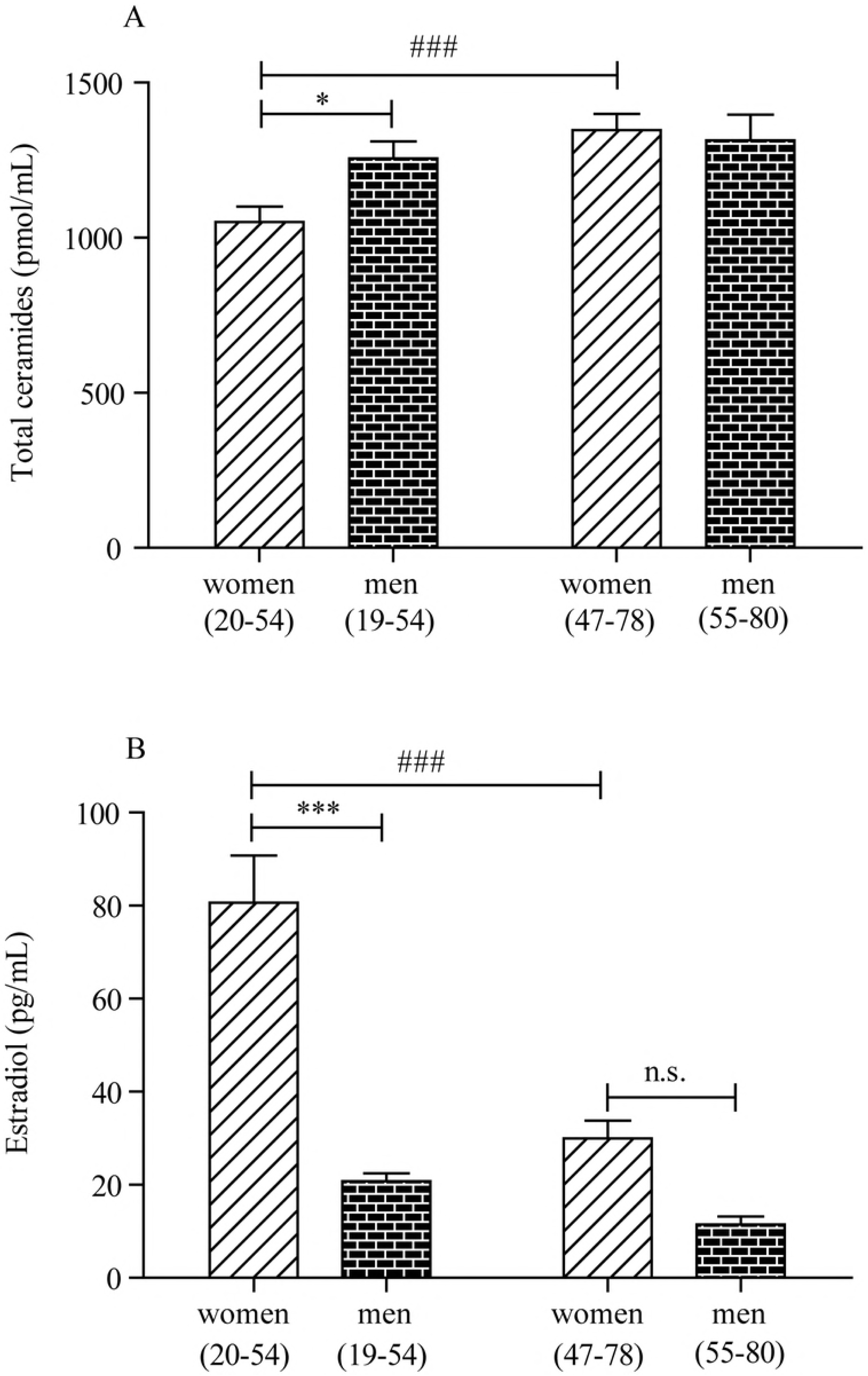
Plasma ceramide and estradiol concentrations in men and women. (A) Plasma ceramide levels in, left, pre-menopausal women (20-54 years, n = 44) and age-matched men (19-54 years, n = 48) and, right, post-menopausal women (47-78 years, n = 40) and age-matched men (55-80 years, n = 32). (B) Plasma estradiol levels in, left, pre-menopausal women (20-54 years, n = 44) and age-matched men (19-54 years, n = 48) and, right, post-menopausal women (47-78 years, n = 40) and age-matched men (55-80 years, n = 32). ^∗^*p* < .05, ^∗∗^*p* < .01, ^∗∗∗^*p* < .001; 2-way ANOVA followed by Bonferroni post-hoc test (women 20-54 years versus men 19-54 years). # *p* < .05, ## *p* < .01, ### *p* < .001; 2-way ANOVA followed by Bonferroni post-hoc test (women 20-54 years versus women 47-78 years).

### Variables associated with plasma ceramides

Next, we used multivariable linear regression models to test the association between age and ceramides and adjust for potential covariates for which data had been collected (Table 1). These factors included hypertension (32/164 subjects, 16 women), tobacco smoking (23/164 subjects, 17 women), use of anti-hypercholesterolemic (12/164 subjects, 3 women) or contraceptive agents (6/164 subjects, 6 women), obesity (0/164 subjects) and diabetes (0/164 subjects). We did not take into account the number of cigarettes smoked as a variable of multivariable linear regression analysis. The adjusted linear regression analysis confirmed that ceramide (d18:1/24:0) (β (SE) = 5.67 (2.38); *p* = .0198) and ceramide (d18:1/24:1) (β (SE) = 2.88 (0.61); *p* < .0001) were positively associated with age in women and also, unexpectedly, revealed an opposite, albeit weaker, trend with ceramide (d18:1/16:0) (β (SE) = -.08 (.04); *p* = .0285) (Table 2A).

**Table 2A.**
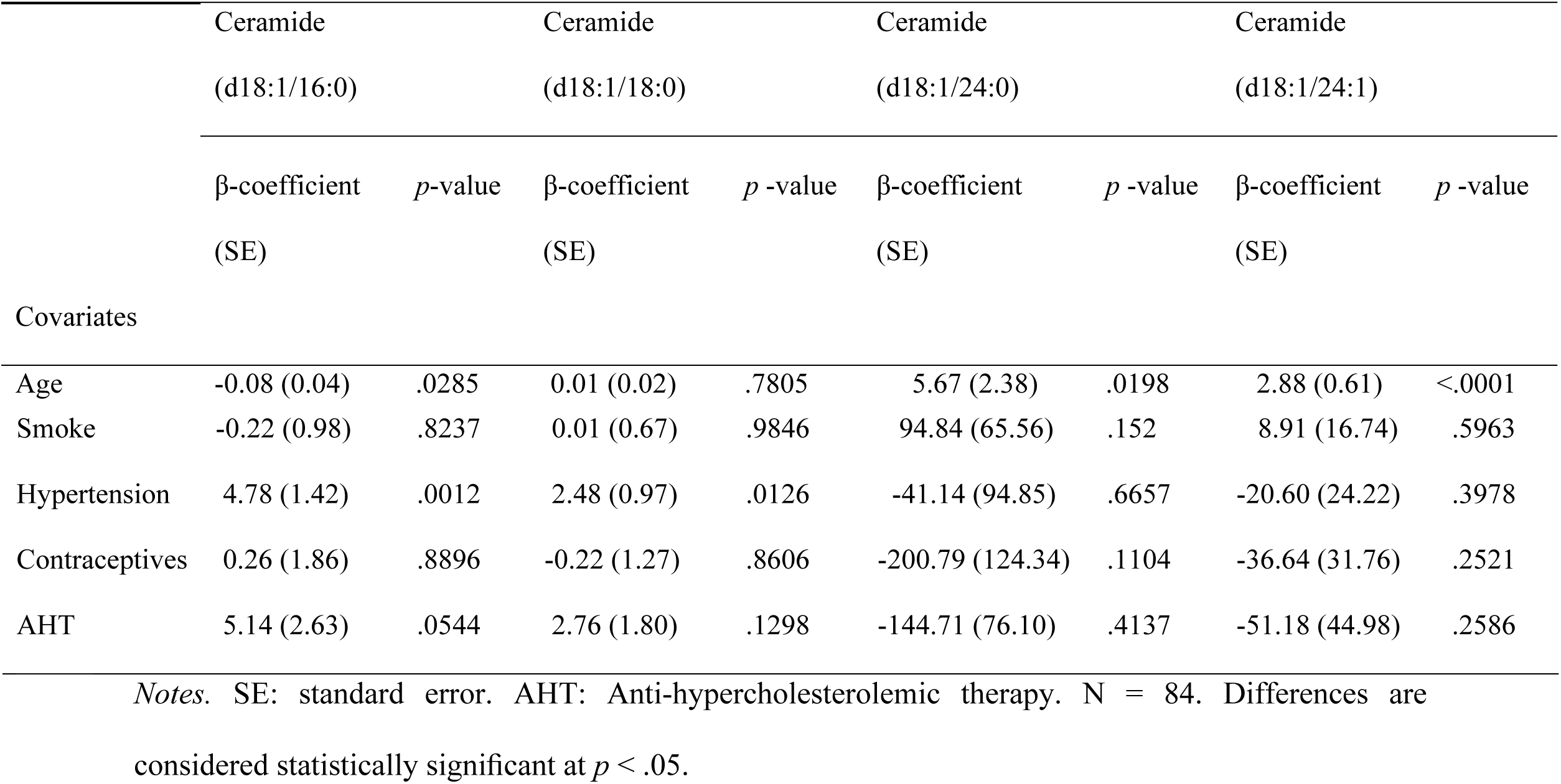
Multivariable linear regression analysis to assess the association between individual ceramide species and variables (age, smoke, hypertension, contraceptive use, anti-hypercholesterolemic therapy) among women.

|  | Ceramide<br>(d18:1/16:0) |  | Ceramide<br>(d18:1/18:0) |  | Ceramide<br>(d18:1/24:0) |  | Ceramide<br>(d18:1/24:1) |  |
| --- | --- | --- | --- | --- | --- | --- | --- | --- |
| | β-coefficient<br>(SE) | $p$-value | β-coefficient<br>(SE) | $p$-value | β-coefficient<br>(SE) | $p$-value | β-coefficient<br>(SE) | $p$-value |
| Covariates |  |  |  |  |  |  |  |  |
| Age | -0.08 (0.04) | .0285 | 0.01 (0.02) | .7805 | 5.67 (2.38) | .0198 | 2.88 (0.61) | <.0001 |
| Smoke | -0.22 (0.98) | .8237 | 0.01 (0.67) | .9846 | 94.84 (65.56) | .152 | 8.91 (16.74) | .5963 |
| Hypertension | 4.78 (1.42) | .0012 | 2.48 (0.97) | .0126 | -41.14 (94.85) | .6657 | -20.60 (24.22) | .3978 |
| Contraceptives | 0.26 (1.86) | .8896 | -0.22 (1.27) | .8606 | -200.79 (124.34) | .1104 | -36.64 (31.76) | .2521 |
| AHT | 5.14 (2.63) | .0544 | 2.76 (1.80) | .1298 | -144.71 (76.10) | .4137 | -51.18 (44.98) | .2586 |

Notes. SE: standard error. AHT: Anti-hypercholesterolemic therapy. N = 84. Differences are considered statistically significant at $p < .05$ .

In men (Table 2B), the analysis unmasked a statistically detectable association between age and ceramide (d18:1/24:1) (β (SE) = 1.86 (.77); *p* = .0179).

**Table 2B.**
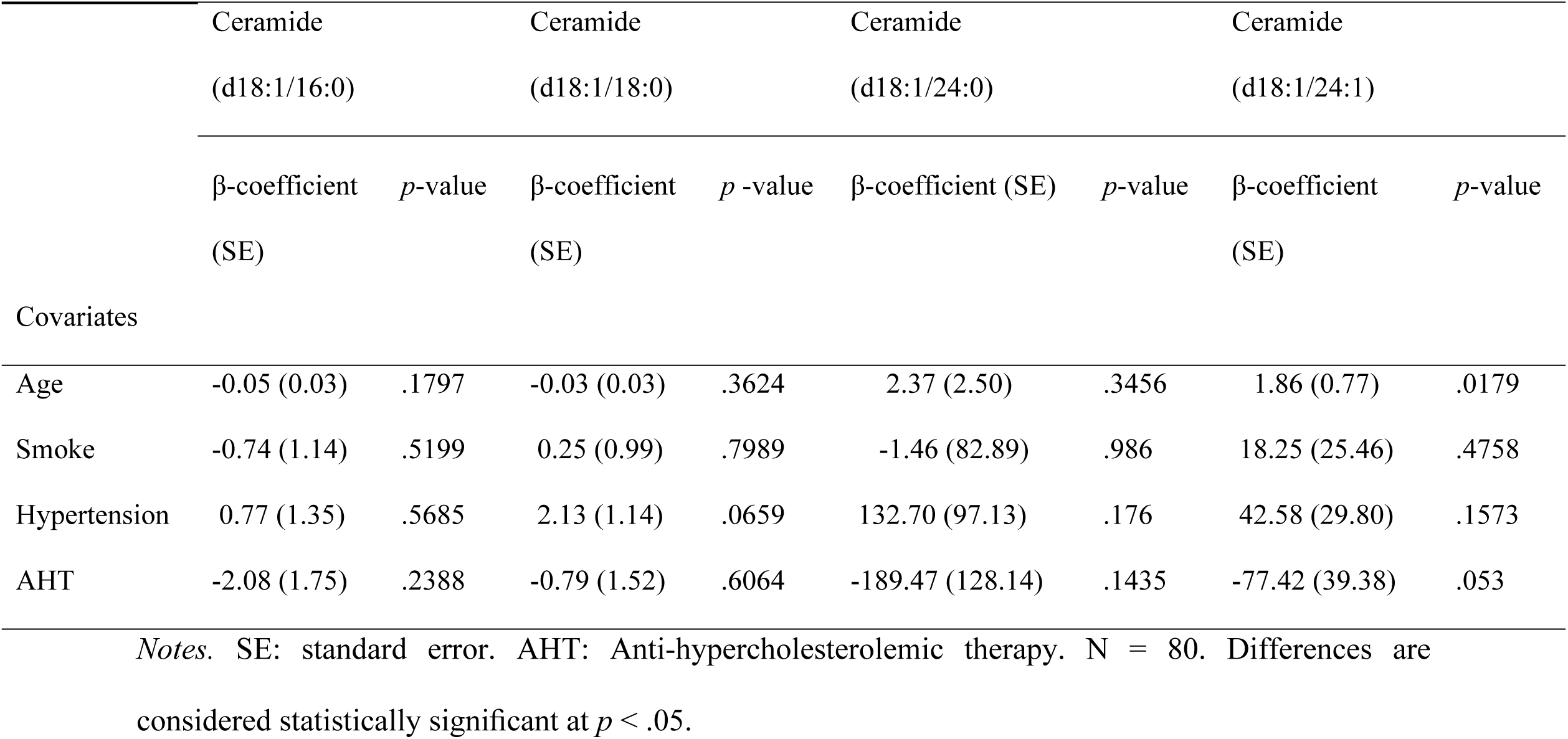
Multivariable linear regression analysis to assess the association between individual ceramide species and variables (age, smoke, hypertension, contraceptive use, anti-hypercholesterolemic therapy) among men.

|  | Ceramide<br>(d18:1/16:0) |  | Ceramide<br>(d18:1/18:0) |  | Ceramide<br>(d18:1/24:0) |  | Ceramide<br>(d18:1/24:1) |  |
| --- | --- | --- | --- | --- | --- | --- | --- | --- |
| | β-coefficient<br>(SE) | $p$-value | β-coefficient<br>(SE) | $p$-value | β-coefficient (SE) | $p$-value | β-coefficient<br>(SE) | $p$-value |
| Covariates |  |  |  |  |  |  |  |  |
| Age | -0.05 (0.03) | .1797 | -0.03 (0.03) | .3624 | 2.37 (2.50) | .3456 | 1.86 (0.77) | .0179 |
| Smoke | -0.74 (1.14) | .5199 | 0.25 (0.99) | .7989 | -1.46 (82.89) | .986 | 18.25 (25.46) | .4758 |
| Hypertension | 0.77 (1.35) | .5685 | 2.13 (1.14) | .0659 | 132.70 (97.13) | .176 | 42.58 (29.80) | .1573 |
| AHT | -2.08 (1.75) | .2388 | -0.79 (1.52) | .6064 | -189.47 (128.14) | .1435 | -77.42 (39.38) | .053 |

Notes. SE: standard error. AHT: Anti-hypercholesterolemic therapy. N = 80. Differences are considered statistically significant at $p < .05$ .

Interestingly, the analysis also pointed to a significant association, found only in women, between hypertension and ceramide (d18:1/16:0) (β (SE) = 4.78 (1.42); *p* = .0012) and (d18:1/18:0) (β (SE) = 2.48 (.97); *p* = .0126) (Table 2A). Of note, 15 out of 16 women affected by hypertension were in the post-menopausal group. Finally, we found no associations, in either sex, between ceramide levels and any other variable, including smoking, contraceptive use or anti-hypercholesterolemic agents, obesity and diabetes (Table 2A-B).

### Plasma ceramide levels are negatively correlated with estradiol in women, but not in men

Next, we investigated a possible correlation between plasma levels of estradiol, which are known to fall significantly at menopause, and ceramides. Estradiol was measured with a competitive binding immunoassay. As expected, plasma estradiol was higher in pre-menopausal women (<55 years) compared to men of the same age (*p* < .001, 2-way ANOVA followed by Bonferroni post-hoc test) (Fig 3B). After menopause, estradiol levels sharply decreased in both sexes (Fig 3B). Fig 4 illustrates the results of Pearson’s analyses of ceramide levels in female subjects of all ages. The results show a statistically significant negative correlation between estradiol and ceramide (d18:1/24:1) (r = −0.294; *p* = .007), a non-significant negative trend between estradiol and ceramide (d18:1/24:0) (r = −0.202; *p* = .066) and no correlations between estradiol and other ceramide species.

**Fig 4.**
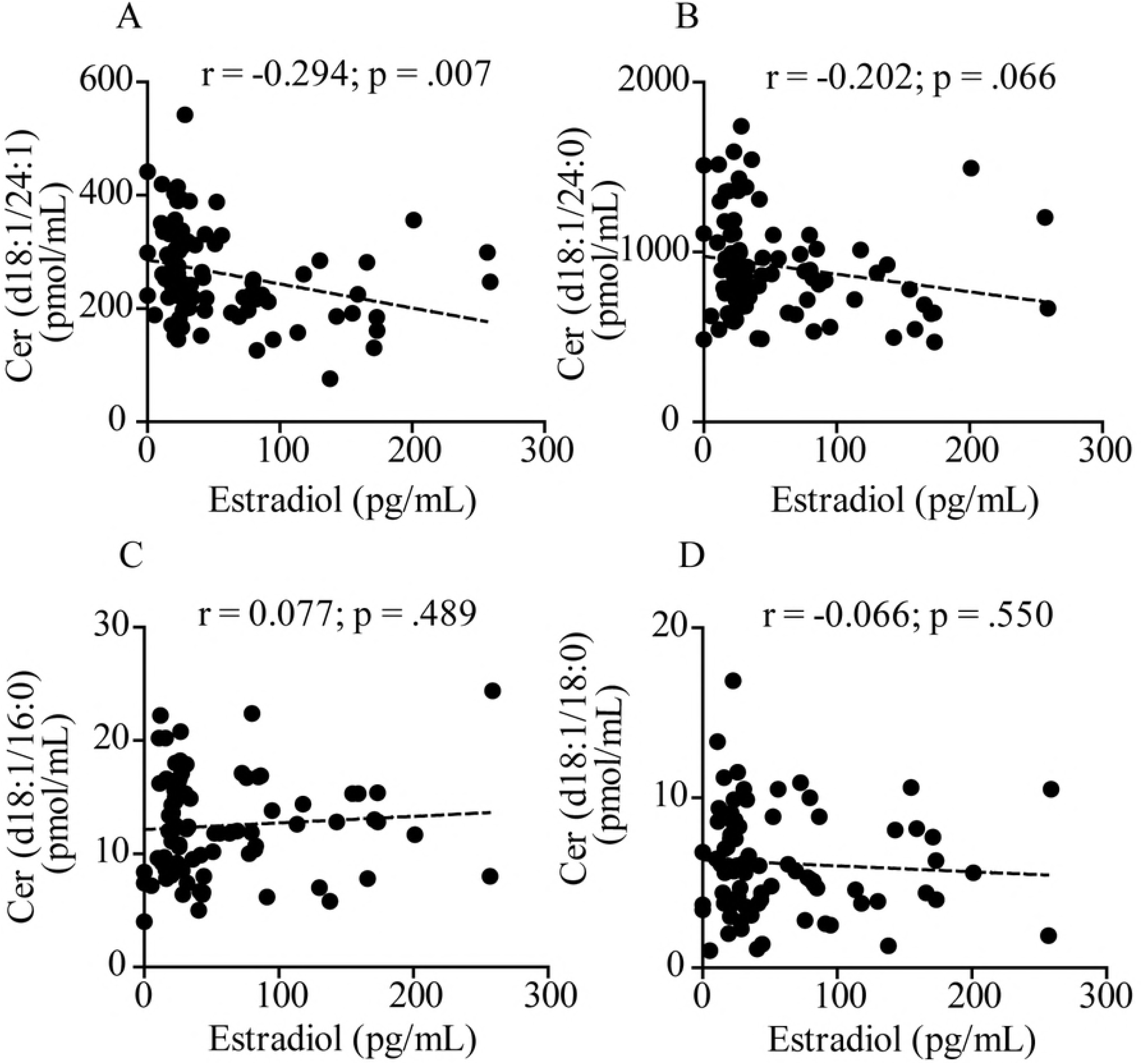
Pearson’s correlation analysis between estradiol and levels of various ceramide species in plasma from 84 women aged 20 to 78 years. (A) Ceramide (d18:1/24:1); (B) Ceramide (d18:1/24:0); (C) Ceramide (d18:1/16:0); (D) Ceramide (d18:1/18:0). Correlation is considered statistically significant at *p* < .05.

By contrast, in men, no correlation was observed between estradiol and any ceramide species, including ceramide (d18:1/24:1) (r = −0.034; *p* = .763) (Fig 5), which was found to be correlated with aging (β (SE) = 1.86 (.77); *p* = .0179) (Table 2B).

**Fig 5.**
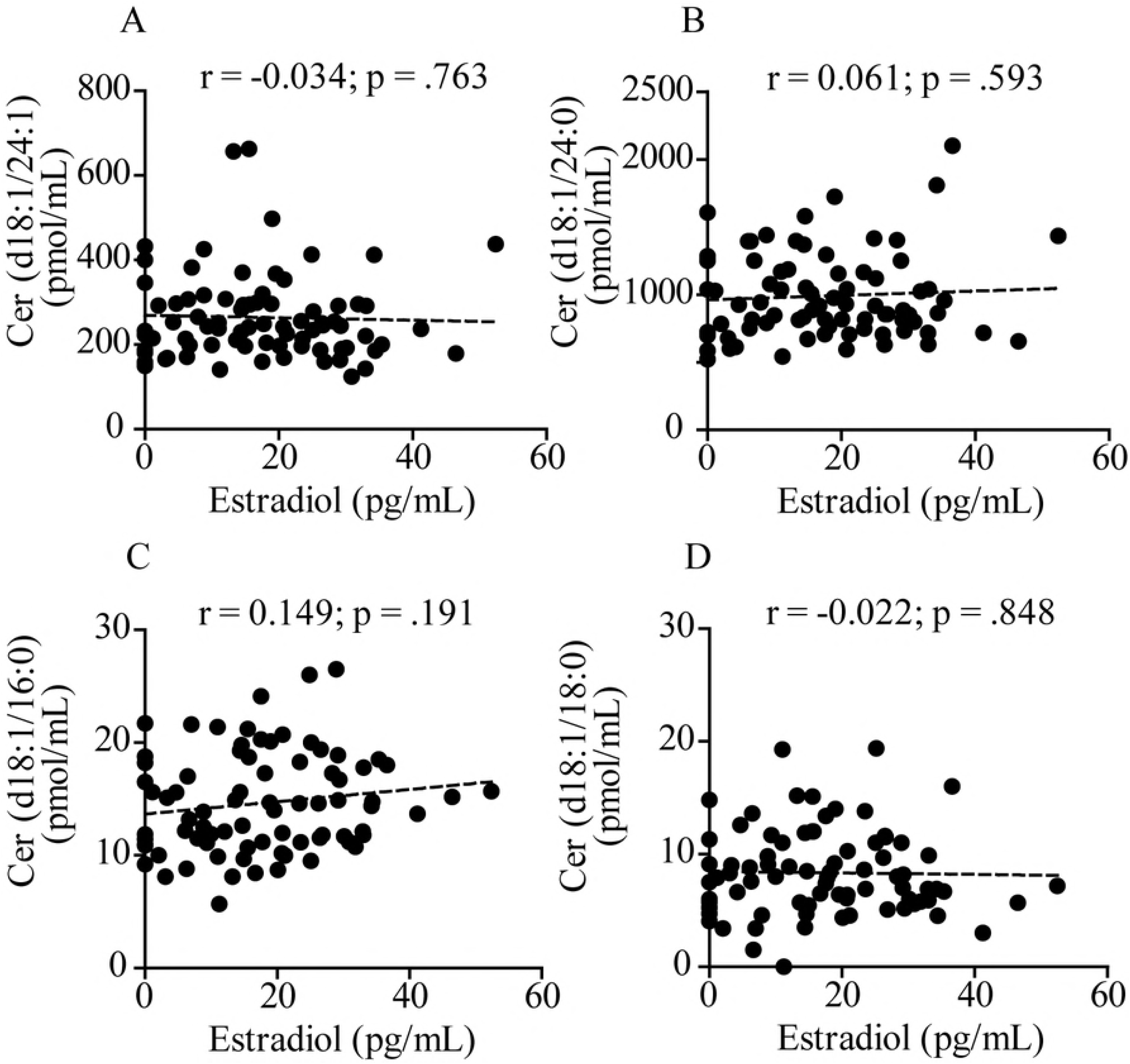
Pearson’s correlation analysis between estradiol and level of various ceramide species in plasma from 80 men aged 19 to 80 years. (A) Ceramide (d18:1/24:1); (B) Ceramide (d18:1/24:0); (C) Ceramide (d18:1/16:0); (D) Ceramide (d18:1/18:0). Correlation is considered statistically significant at *p* < .05.

### Estradiol suppresses ceramide accumulation *in vitro*

Sphingolipid-derived mediators regulate steroidogenesis [32], but it is still unknown whether estrogen hormones influence sphingolipid metabolism. To gain insight into the causality of the negative correlation observed between estradiol and ceramide in women, we asked whether the sex hormone might regulate ceramide mobilization (i.e. formation and/or degradation) in human MCF7 breast cancer cells, a cell line that expresses the estrogen receptor α (ERα) and β (ERβ) [31]. MCF7 cells were treated with estradiol (10 nM) for 24 h and ceramides quantified in lipid extracts by LC-MS/MS. Results indicate that the exposure to estradiol causes a substantial reduction in ceramide (d18:1/16:0) (*p* = .02, unpaired Student’s *t*-test), (d18:1/24:0) (*p* = .0002) and (d18:1/24:1) (*p* = .0006) (Table 3), thereby suggesting that estradiol causes a downward regulation in the mobilization of these ceramides.

**Table 3.** Effects of estradiol on ceramide levels in MCF7 human breast cancer cells. Cells were treated for 24 h with vehicle (0.1% DMSO in phenol-free DMEM) or 17-β-estradiol (10 nM) and ceramide levels were measured by LC-MS/MS.

|  | Vehicle | Estradiol | *p*-value |
| --- | --- | --- | --- |
| Ceramide | (pmol/mg protein) | (pmol/mg protein) |  |
| (d18:1/16:0) | 57.4 ± 6.2 | 41.8 ± 0.8 | .02 |
| (d18:1/18:0) | 16.9 ± 1.3 | 13.3 ± 1.6 | .12 |
| (d18:1/24:0) | 209.4 ± 14.1 | 134.6 ± 7.4 | .0002 |
| (d18:1/24:1) | 265.6 ± 13.4 | 184.6 ± 12.6 | .0006 |

## Discussion

In the present study, we investigated the age- and sex-dependent trajectories of plasma ceramides in 80 men and 84 women aged 19-80 years. The subjects were not affected by any major illness or medical condition known to be linked to alterations in circulating ceramide levels. The results show that, in women, plasma levels of two ceramide species – (d18:1/24:0) and (d18:1/24:1) – increased with age, and that this change cannot be ascribed to confounding factors such as obesity, diabetes, hypertension, tobacco smoking or ongoing anti-cholesterol and contraceptive therapy. In men, the analysis revealed a statistically detectable association between age and ceramide (d18:1/24:1). Further analyses identified a significant negative correlation between circulating levels of estradiol and ceramide (d18:1/24:1) in women of all ages, but not in men. Finally, *in vitro* experiments in a human cell line expressing estrogen ER-α and ER-β receptors showed that treatment with exogenous estradiol produced a significant decrease in ceramide accumulation. These findings suggest that aging is accompanied, in humans, by an increase in the plasma concentrations of two ceramide species, (d18:1/24:0) and (d18:1/24:1), which are also known to be elevated in age-dependent pathologies, such as atherosclerosis and cardiovascular disease [33,34]. The results also point to the intriguing possibility that estradiol might control circulating ceramide levels in a sexually dimorphic manner.

Previous studies have reported sex-dependent differences in plasma ceramides, but with somewhat contrasting results. In a small investigation of blood serum samples from 10 Caucasian volunteers (5 males aged 27-33 years and 5 females aged 26-33 years), ceramide (42:1) was found to be higher in women compared to men [16]. Another study, performed on a much larger cohort of young Mexican Americans (1,076 individuals, 39.1% males, median age 35.7 years) [17], uncovered an association between plasma ceramides and sex after adjusting for age and body mass index: ceramide levels were lower in women than in men. These disparities were mostly driven by long-chain ceramide species, such as (d18:1/22:0), (d18:1/24:0) and (d18:1/24:1). Finally, in a multiethnic population sample of 366 women and 626 men aged over 55 years, enrolled in the Baltimore Longitudinal Study of Aging, plasma ceramide concentrations were found to be higher in women compared to men [18].

These studies did not focus on age as a variable. By contrast, in the present work we set out to address the impact of aging on ceramide levels and excluded subjects with disease conditions that had been previously shown to affect ceramides, such as diabetes [35], cancer [36], renal disease [37], cardiovascular disease [38], and obesity [39]. We did not exclude subjects with hypertension, however, because the association of this disease state with altered ceramides remains to be fully established [11,18]. In our sample, we were able to confirm the presence of age- and sex-dependent differences in the plasma concentrations of certain ceramide species, but not others. A multivariable analysis showed that, in women, aging is accompanied by increased levels of ceramide (d18:1/24:0) and (d18:1/24:1). In men, the analysis also unmasked a statistically detectable association between age and ceramide (d18:1/24:1). These age-dependent changes could not be ascribed to obesity, diabetes, tobacco cigarette smoking and use of anti-hypercholesterolemic or contraceptive agents. Interestingly, secondary data analyses found that ceramide levels were significantly lower in female than male subjects aged 20-54, a difference that disappeared after menopause. In their 2015 study, Mielke and collaborators did not specifically include menopause as a variable but suggested that menopause and estradiol may influence ceramide levels [18]. Two sets of results presented here support this prediction. First, we showed that, in women of all ages, plasma estradiol was negatively correlated with ceramide (d18:1/24:1) and displayed a trend toward correlation with ceramide (d18:1/24:0). No such relationship was detectable in men. Second, we found that incubation with exogenous estradiol (10 nM) lowered the levels of various ceramides, including ceramide (d18:1/24:0) and (d18:1/24:1), in human estrogen-sensitive MCF7 cells. The findings outlined above suggest that age-dependent changes in estradiol may affect ceramide metabolism differentially in men and women. The correlation between estradiol and ceramide raises the possibility that changes in the availability of certain ceramide species [e.g. ceramide (d18:1/24:1)] might be implicated in the reported cardioprotective, antihypertensive and neuroprotective effects of estradiol [40-42]. In this context, it is important to point out that increased ceramide levels have been consistently linked to heightened risk of myocardial infarction and stroke [43]. Moreover, recent evidence indicates that alterations of ceramide levels may mirror, indirectly, ongoing neurodegenerative processes and might be used as a biomarker for Alzheimer’s disease development and progression [44]. Interestingly, the close association between alterations in ceramide levels and the likelihood of developing cognitive impairment and Alzheimer-related pathology may be particularly strong in women, thereby indicating that sex-related mechanisms might participate in shaping the Alzheimer phenotype. Of note, elderly women show changes in plasma ceramide levels at the onset of their memory impairment [15]. In the past few years, the neurobiology of memory disorders has received increasing attention [45]. Memory deficits are often reported by women in temporal proximity of menopause [46], and recent findings indicate distinct changes in memory processing that appear to be linked more to the pre-menopausal status than to chronological aging: in peri-menopausal women, the onset and progression of cognitive decline are often associated with a menopause-related decrease in estradiol levels [47]. Thus, the interplay between estradiol and ceramides, described here, may potentially be of broad significance for a variety of age-dependent disorders, including cardiovascular disease and cognitive impairment or any intersection of the two conditions [48].

The present study has several limitations. First, even though we excluded persons affected by obesity and diabetes, we did not collect information on metabolic factors that could potentially impact ceramide mobilization – such as adiposity, physical activity and levels of cholesterol and glucose in blood [18,49-51]. Second, our study was focused on a specific subset of ceramides that we had previously found to be altered in the hippocampus of aged male and female mice [19]. This group of ceramides has been proposed as potential biomarker for cardiovascular risk [52], but is still only a small fraction of the vast number of ceramides produced by the body. Third, we measured total circulating levels of ceramides and did not attempt to separate ceramide pools bound to specific plasma lipoproteins [53]. Because the distribution of ceramides among lipoproteins may change with obesity and diabetes [50], it is possible that aging might exert a similar effect.

Our findings also raise a number of relevant questions, which should be addressed in future work. First, even though the results suggest that estradiol modulates ceramide mobilization in women, the precise mechanism and functional significance of this effect remain to be determined. One possibility is that activation of estrogen receptors results in the down-regulation of *de novo* ceramide biosynthesis, for example by suppressing the expression of key enzymes such as serine-palmitoyltransferase and ceramide synthase [54]. Alternatively, estradiol might stimulate the expression of ceramide-hydrolyzing enzymes such as acid or neutral ceramidase. Probing the link between estradiol and ceramides will require additional experimentation, which may include measuring circulating ceramide levels in women throughout the menstrual cycle or assessing the impact of endogenous and exogenous estradiol on sphingolipid metabolism in female animals. At the functional level, studies are needed to correlate ceramide and estrogen levels to a broad panel of biomarkers (e.g., lipoprotein profile, C-reactive protein) and clinical outcome measures (e.g., future adverse cardiovascular events and cognitive impairment) [52]. Without such information, the clinical significance of our findings remains speculative. Second, the lack of correlation between estradiol and ceramide (d18:1/18:0) and lack of association with age implies that this ceramide species, though elevated after menopause, may be subjected to a different regulation than ceramide (d18:1/24:1), whose levels are correlated with estradiol and are statistically associated with age. Third, we observed a substantial age-dependent decrease in plasma dihydroceramide (d18:0/24:0) in men aged 54-80 years. The significance of this finding is presently unclear, but warrants further attention. Fourth, we unexpectedly uncovered an association between hypertension in post-menopausal women and elevated levels of ceramide (d18:1/16:0) and (d18:1/18:0). While this result is consistent with previous reports [11], caution is warranted until studies with a larger cohort of pre- and post-menopausal women are performed. Finally, as mentioned above, the relation between ceramide levels and premorbid changes in cardiovascular, metabolic or cognitive function was not investigated in our study and deserves further investigations. Despite these unanswered questions, our results reveal the existence of a link between age, estradiol, and ceramides, which might contribute to age-dependent pathologies in post-menopausal women.

## References

1. Castro BM, Prieto M, Silva LC (2014) Ceramide: a simple sphingolipid with unique biophysical properties. Prog Lipid Res 54: 53–67.

2. Grassme H, Riethmuller J, Gulbins E (2007) Biological aspects of ceramide-enriched membrane domains. Prog Lipid Res 46: 161–170.

3. Garcia-Barros M, Coant N, Kawamori T, Wada M, Snider AJ, et al. (2016) Role of neutral ceramidase in colon cancer. FASEB J 30: 4159–4171.

4. Saddoughi SA, Ogretmen B (2013) Diverse functions of ceramide in cancer cell death and proliferation. Adv Cancer Res 117: 37–58.

5. Bieberich E (2012) Ceramide and sphingosine-1-phosphate signaling in embryonic stem cell differentiation. Methods Mol Biol 874: 177–192.

6. Venable ME, Lee JY, Smyth MJ, Bielawska A, Obeid LM (1995) Role of ceramide in cellular senescence. J Biol Chem 270: 30701–30708.

7. Morad SA, Cabot MC (2013) Ceramide-orchestrated signalling in cancer cells. Nat Rev Cancer 13: 51–65.

8. Maeng HJ, Song JH, Kim GT, Song YJ, Lee K, et al. (2017) Celecoxib-mediated activation of endoplasmic reticulum stress induces de novo ceramide biosynthesis and apoptosis in hepatoma HepG2 cells mobilization. BMB Rep 50: 144–149.

9. Samad F, Hester KD, Yang G, Hannun YA, Bielawski J (2006) Altered adipose and plasma sphingolipid metabolism in obesity: a potential mechanism for cardiovascular and metabolic risk. Diabetes 55: 2579–2587.

10. Haus JM, Kashyap SR, Kasumov T, Zhang R, Kelly KR, et al. (2009) Plasma ceramides are elevated in obese subjects with type 2 diabetes and correlate with the severity of insulin resistance. Diabetes 58: 337–343.

11. Spijkers LJ, van den Akker RF, Janssen BJ, Debets JJ, De Mey JG, et al. (2011) Hypertension is associated with marked alterations in sphingolipid biology: a potential role for ceramide. PLoS One 6: e21817.

12. Ichi I, Nakahara K, Miyashita Y, Hidaka A, Kutsukake S, et al. (2006) Association of ceramides in human plasma with risk factors of atherosclerosis. Lipids 41: 859–863.

13. de Mello VD, Lankinen M, Schwab U, Kolehmainen M, Lehto S, et al. (2009) Link between plasma ceramides, inflammation and insulin resistance: association with serum IL-6 concentration in patients with coronary heart disease. Diabetologia 52: 2612–2615.

14. Mielke MM, Bandaru VV, Haughey NJ, Rabins PV, Lyketsos CG, et al. (2010) Serum sphingomyelins and ceramides are early predictors of memory impairment. Neurobiol Aging 31: 17–24.

15. Mielke MM, Haughey NJ, Bandaru VV, Schech S, Carrick R, et al. (2010) Plasma ceramides are altered in mild cognitive impairment and predict cognitive decline and hippocampal volume loss. Alzheimers Dement 6: 378–385.

16. Ishikawa M, Tajima Y, Murayama M, Senoo Y, Maekawa K, et al. (2013) Plasma and serum from nonfasting men and women differ in their lipidomic profiles. Biol Pharm Bull 36: 682–685.

17. Weir JM, Wong G, Barlow CK, Greeve MA, Kowalczyk A, et al. (2013) Plasma lipid profiling in a large population-based cohort. J Lipid Res 54: 2898–2908.

18. Mielke MM, Bandaru VV, Han D, An Y, Resnick SM, et al. (2015) Demographic and clinical variables affecting mid- to late-life trajectories of plasma ceramide and dihydroceramide species. Aging Cell 14: 1014–1023.

19. Vozella V, Basit A, Misto A, Piomelli D (2017) Age-dependent changes in nervonic acid-containing sphingolipids in mouse hippocampus. Biochim Biophys Acta 1862: 1502–1511.

20. Green CL, Mitchell SE, Derous D, Wang Y, Chen L, et al. (2017) The effects of graded levels of calorie restriction: IX. Global metabolomic screen reveals modulation of carnitines, sphingolipids and bile acids in the liver of C57BL/6 mice. Aging Cell 16: 529–540.

21. Folstein MF, Folstein SE, McHugh PR (1975) “Mini-mental state”. A practical method for grading the cognitive state of patients for the clinician. J Psychiatr Res 12: 189–198.

22. Carlesimo GA, Caltagirone C, Gainotti G (1996) The Mental Deterioration Battery: normative data, diagnostic reliability and qualitative analyses of cognitive impairment. The Group for the Standardization of the Mental Deterioration Battery. Eur Neurol 36: 378–384.

23. McKhann GM, Knopman DS, Chertkow H, Hyman BT, Jack CR, Jr., et al. (2011) The diagnosis of dementia due to Alzheimer’s disease: recommendations from the National Institute on Aging-Alzheimer’s Association workgroups on diagnostic guidelines for Alzheimer’s disease. Alzheimers Dement 7: 263–269.

24. Petersen RC, Morris JC (2005) Mild cognitive impairment as a clinical entity and treatment target. Arch Neurol 62: 1160–1163; discussion 1167.

25. First MB, Pincus HA (2002) The DSM-IV Text Revision: rationale and potential impact on clinical practice. Psychiatr Serv 53: 288–292.

26. Mencarelli C, Martinez-Martinez P (2013) Ceramide function in the brain: when a slight tilt is enough. Cell Mol Life Sci 70: 181–203.

27. Filippov V, Song MA, Zhang K, Vinters HV, Tung S, et al. (2012) Increased ceramide in brains with Alzheimer’s and other neurodegenerative diseases. J Alzheimers Dis 29: 537–547.

28. Iorio M, Spalletta G, Chiapponi C, Luccichenti G, Cacciari C, et al. (2013) White matter hyperintensities segmentation: a new semi-automated method. Front Aging Neurosci 5: 76.

29. Basit A, Piomelli D, Armirotti A (2015) Rapid evaluation of 25 key sphingolipids and phosphosphingolipids in human plasma by LC-MS/MS. Anal Bioanal Chem 407: 5189–5198.

30. Soule HD, Vazguez J, Long A, Albert S, Brennan M (1973) A human cell line from a pleural effusion derived from a breast carcinoma. J Natl Cancer Inst 51: 1409–1416.

31. Brooks SC, Locke ER, Soule HD (1973) Estrogen receptor in a human cell line (MCF-7) from breast carcinoma. J Biol Chem 248: 6251–6253.

32. Lucki NC, Sewer MB (2010) The interplay between bioactive sphingolipids and steroid hormones. Steroids 75: 390–399.

33. Pan W, Yu J, Shi R, Yan L, Yang T, et al. (2014) Elevation of ceramide and activation of secretory acid sphingomyelinase in patients with acute coronary syndromes. Coron Artery Dis 25: 230–235.

34. Yu J, Pan W, Shi R, Yang T, Li Y, et al. (2015) Ceramide is upregulated and associated with mortality in patients with chronic heart failure. Can J Cardiol 31: 357–363.

35. Galadari S, Rahman A, Pallichankandy S, Galadari A, Thayyullathil F (2013) Role of ceramide in diabetes mellitus: evidence and mechanisms. Lipids Health Dis 12: 98.

36. Kizhakkayil J, Thayyullathil F, Chathoth S, Hago A, Patel M, et al. (2012) Glutathione regulates caspase-dependent ceramide production and curcumin-induced apoptosis in human leukemic cells. Free Radic Biol Med 52: 1854–1864.

37. Mitsnefes M, Scherer PE, Friedman LA, Gordillo R, Furth S, et al. (2014) Ceramides and cardiac function in children with chronic kidney disease. Pediatr Nephrol 29: 415–422.

38. Alewijnse AE, Peters SL (2008) Sphingolipid signalling in the cardiovascular system: good, bad or both? Eur J Pharmacol 585: 292–302.

39. Samad F, Badeanlou L, Shah C, Yang G (2011) Adipose tissue and ceramide biosynthesis in the pathogenesis of obesity. Adv Exp Med Biol 721: 67–86.

40. Lagranha CJ, Silva TLA, Silva SCA, Braz GRF, da Silva AI, et al. (2018) Protective effects of estrogen against cardiovascular disease mediated via oxidative stress in the brain. Life Sci 192: 190–198.

41. Hernandez-Fonseca K, Massieu L, Garcia de la Cadena S, Guzman C, Camacho-Arroyo I (2012) Neuroprotective role of estradiol against neuronal death induced by glucose deprivation in cultured rat hippocampal neurons. Neuroendocrinology 96: 41–50.

42. Mercuro G, Zoncu S, Piano D, Pilia I, Lao A, et al. (1998) Estradiol-17beta reduces blood pressure and restores the normal amplitude of the circadian blood pressure rhythm in postmenopausal hypertension. Am J Hypertens 11: 909–913.

43. Park JY, Lee SH, Shin MJ, Hwang GS (2015) Alteration in metabolic signature and lipid metabolism in patients with angina pectoris and myocardial infarction. PLoS One 10: e0135228.

44. Mielke MM, Lyketsos CG (2006) Lipids and the pathogenesis of Alzheimer’s disease: is there a link? Int Rev Psychiatry 18: 173–186.

45. Jessen F (2014) Subjective and objective cognitive decline at the pre-dementia stage of Alzheimer’s disease. Eur Arch Psychiatry Clin Neurosci 264 Suppl 1: S3–7.

46. Newhouse P, Dumas J (2015) Estrogen-cholinergic interactions: Implications for cognitive aging. Horm Behav 74: 173–185.

47. Henderson VW (2010) Action of estrogens in the aging brain: dementia and cognitive aging. Biochim Biophys Acta 1800: 1077–1083.

48. Toledo JB, Arnold SE, Raible K, Brettschneider J, Xie SX, et al. (2013) Contribution of cerebrovascular disease in autopsy confirmed neurodegenerative disease cases in the National Alzheimer’s Coordinating Centre. Brain 136: 2697–2706.

49. Fabbri E, Yang A, Simonsick EM, Chia CW, Zoli M, et al. (2016) Circulating ceramides are inversely associated with cardiorespiratory fitness in participants aged 54-96 years from the Baltimore Longitudinal Study of Aging. Aging Cell 15: 825–831.

50. Boon J, Hoy AJ, Stark R, Brown RD, Meex RC, et al. (2013) Ceramides contained in LDL are elevated in type 2 diabetes and promote inflammation and skeletal muscle insulin resistance. Diabetes 62: 401–410.

51. Bergman BC, Brozinick JT, Strauss A, Bacon S, Kerege A, et al. (2015) Serum sphingolipids: relationships to insulin sensitivity and changes with exercise in humans. Am J Physiol Endocrinol Metab 309: E398–408.

52. Summers SA (2017) Could Ceramides Become the New Cholesterol? Cell Metab.

53. Hammad SM, Pierce JS, Soodavar F, Smith KJ, Al Gadban MM, et al. (2010) Blood sphingolipidomics in healthy humans: impact of sample collection methodology. J Lipid Res 51: 3074–3087.

54. Gault CR, Obeid LM, Hannun YA (2010) An overview of sphingolipid metabolism: from synthesis to breakdown. Adv Exp Med Biol 688: 1–23.

